# ToR-ORd-dynCl: an update of the ToR-ORd model of human ventricular cardiomyocyte with dynamic intracellular chloride

**DOI:** 10.1101/2020.06.01.127043

**Authors:** Jakub Tomek, Alfonso Bueno-Orovio, Blanca Rodriguez

## Abstract

Recently, our group published a new model of human ventricular cardiomyocyte named ToR-ORd (Tomek et al., 2019). Its development, calibration, and validation, were performed using a broad range of human experimental data and brought general insights into modelling of ionic channels. Model calibration ensured the reproduction of key physiological cellular features, with independent multiscale validation demonstrating a correct response to channel blocking drugs and pathophysiological remodelling.

However, for very long simulations (several hours rather than minutes), the ToR-ORd simulations display a drift in its behaviour, caused by modelling chloride concentrations as constant values. This may be a limitation for simulations considering extremely long protocols, or for studies on model stability. To remedy this, we present here an updated version, termed ToR-ORd-dynCl, with dynamic representation of intracellular chloride. This model behaves very similarly to the original ToR-ORd, but with stable properties over long simulations and only a small increase in model complexity.

## 1 Introduction

In a recent study, we presented the development, calibration and independent validation of a human-based ventricular cellular model ToR-ORd (Tomek et al., 2019). Simulations of electrophysiology and excitation-contraction coupling, and their comparison to experimental and clinical data were presented from ionic to whole-organ dynamics, including the electrocardiogram. The ToR-ORd model follows the general structure of previous cardiac cellular models, and specifically the ORd model (O’Hara et al., 2011). Ionic currents and fluxes carrying potassium, sodium, calcium and chloride ions are represented in cell compartments, described by Hodgkin-Huxley equations or Markov models. Intracellular concentrations for potassium, sodium and calcium are dynamically updated based on these currents and fluxes, through ordinary differential equations. For chloride currents, the ToR-ORd model was developed as in Grandi et al. (2010), with chloride concentrations held constant throughout the simulations. This allows to select and maintain specific levels of intracellular chloride, which is an acceptable approximation for simulations of seconds or minutes. However, for very long simulations of the order of several hours, using fixed ionic concentrations (rather than their dynamic update) is known to generate significant drifts in key properties (Hund et al., 2001).

Here we describe an alternative version of the model (named ToR-ORd-dynCl) with dynamic changes in intracellular chloride concentrations. Therefore, in the ToR-ORd-dynCl, intracellular chloride concentrations are not held constant, as in ToR-ORd, but updated dynamically according to its two chloride currents (calcium-sensitive Cl current I_(Ca)Cl_ and background Cl current I_Clb_). We show that simulations with ToR-ORd-dynCl achieve a steady state, and its steady-state behaviour is very similar to ToR-ORd at the time point it was evaluated in (Tomek et al., 2019). The original ToR-ORd is nevertheless suitable for simulations of thousands of beats, whereas for long simulations, the stability of the ToR-ORd-dynCl is a clear advantage. The code for both models is available through https://github.com/jtmff/torord, both for CellML and Matlab.

As this is essentially an addendum, rather than a self-standing article, please also cite the original study (Tomek et al., 2019) whenever referring to this document.

## 2 Methods

Figure 1 illustrates the structure and main features of the ToR-ORd model, highlighting the modifications in the chloride concentrations, introduced in the ToR-ORd-dynCl. The main compartments represented are the main cytosolic space, junctional subspace, and the sarcoplasmic reticulum (SR, further subdivided into junctional and network SR). The following changes to the original ToR-ORd model were implemented:

**Figure 1:**
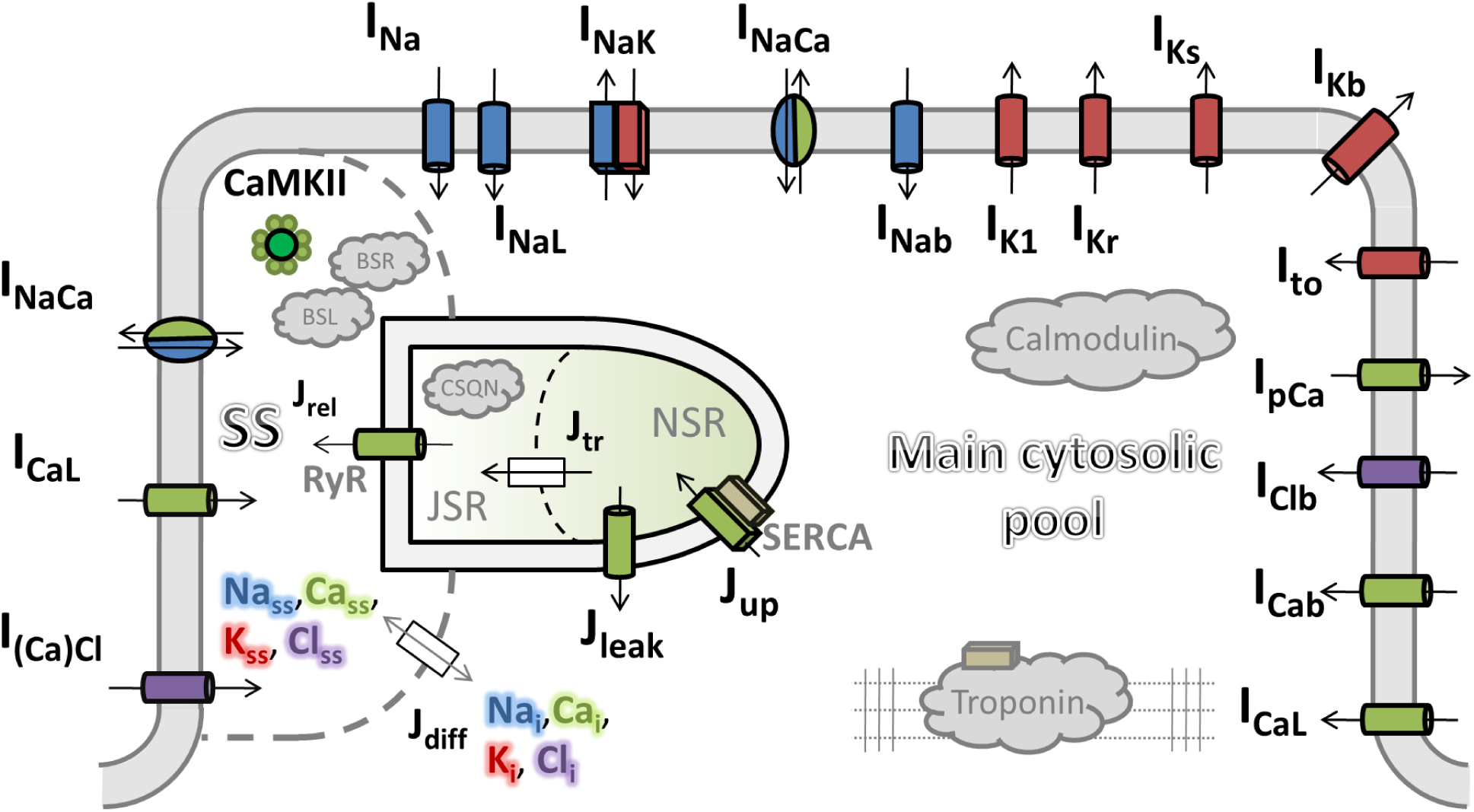
Diagram of ToR-ORd-dynCl. The diagram is based on the of the original ToR-ORd (Tomek et al., 2019), showing model compartments (main cytosolic pool, junctional subspace SS, and subcompartments of the sarcoplasmic reticulum), currents, fluxes, and buffers. Intracellular concentration of ions are listed in the main cytosolic pool and SS compartments using color labels with a halo. Compared to the original model dynamically updating concentrations of sodium, calcium, and potassium, ToR-ORd-dynCl also dynamically updates chloride concentrations.

1. Two state variables were added, corresponding to intracellular concentrations of Cl in myoplasm and junctional subspace.
2. The chloride currents use in their equations the equilibrium potentials based on chloride concentrations in their corresponding compartments.
3. The diffusion between junctional subspace and main cytoplasm was set as:

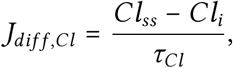

where *Cl*_*ss*_ is the chloride concentration in junctional subspace *Cl_i_* and is the chloride concentration in myoplasm. *τ*_*Cl*_ = 2 ms has the same value as the other two monovalent ions in the model, sodium and potassium.
4. The chloride concentrations in the two subspaces are updated similarly to other ions as follows:

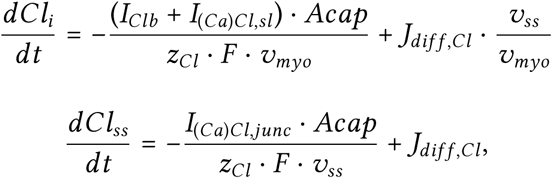

where *v*_*ss*_ and *v*_*myo*_ are the volumes of junctional subspace compartment and myoplasm compartment respectively. *I*_*(Ca)Cl,sl*_ is the Ca-sensitive Cl current in the sarcolemma (linking extracellular and myoplasmic space), whereas *I*_*(Ca)Cl,junc*_ is the Ca-sensitive Cl current in the junctional membrane (linking extracellular space and junctional subspace). We note that in the model, the Ca-sensitive Cl current is wholly placed in the junctional membrane, so *I*_*(Ca)Cl,sl*_ is always zero, but we give the general equation in case future models have a different distribution of the current between sarcolemma and junctional membrane. *z*_*Cl*_ = −1 is the valence of chloride, and *F* = 96485 is the Faraday constant.
5. When the driving force is computed for I_CaL_ in the myoplasm and junctional subspace, it uses the corresponding chloride concentrations in myoplasm and subspace to compute the ionic strength there.
6. In addition, the *τ*_*jca*_ parameter (used in the computation of I_CaL_) was reduced from 75 to 72.5 ms. The minor change was carried out so that the model manifests visible EADs at the same conditions as studied in the original article (0.25 Hz pacing, 85% I_Kr_ block, extracellular Na of 137 mM and extracellular Ca of 2 mM). Without the change in *τ*_*jca*_, the dynamic-chloride model showed only a weak EAD (nevertheless showing a strong EAD if 86% I_Kr_ block was used, demonstrating that only a very minor shift in behaviour was induced by the introduction of dynamic chloride concentration).

## 3 Results and Discussion

### 3.1 Model stability

To visualise long-term stability of ToR-ORd and ToR-ORd-dynCl, we simulated both models for over 27 hours (100 000 beats at 1 Hz), measuring a range of features in the last beat of each thousand. Both models started in the same initial state used for ToR-ORd (with the initial vector of state variables in ToR-ORd-dynCl being extended with *Cl*_*i*_ = *Cl*_*ss*_ = 24 mM).

The action potential duration at 90% repolarisation level (APD90) changes very little in both models throughout the 100 000 beats (Figure 2A), showing small-amplitude oscillations in ToR-ORd-dynCl, and a minor drift in ToR-ORd. However, while ToR-ORd-dynCl similarly stabilises with regards to resting membrane potential, the original ToR-ORd shows a more pronounced drift (Figure 2B). The calcium transient amplitude and diastolic calcium concentration show a moderate elevation in ToR-ORd over time and a stabilisation with ToR-ORd-dynCl (Figure 2C,D). Diastolic potassium concentration stabilises with ToR-ORd-dynCl, but is supraphysiological at the end of 100 000 beats simulation in ToR-ORd (Figure 2E).

**Figure 2:**
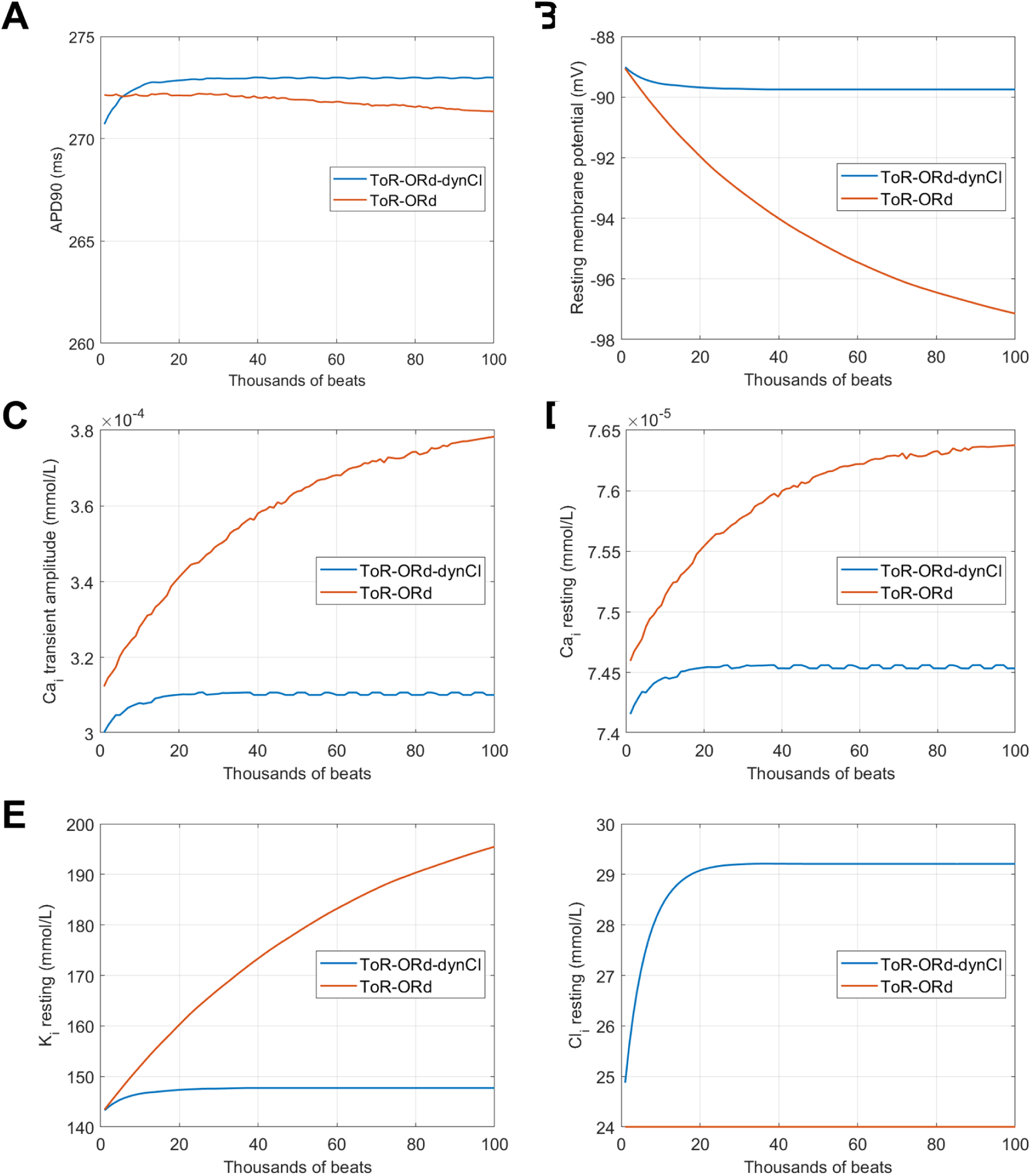
Development of features and selected state variables over time. The plotted values correspond to the feature in the last beat of the given number of thousands of beats. Diastolic ionic concentrations were measured in the main myoplasmic compartment, not in the junctional subspace.

The chloride concentration is held constant in ToR-ORd, and increases to ca. 29.2 mM in ToR-ORd-dynCl (Figure 2F). Based on existing literature, it is difficult to say whether this is physiological or slightly supraphysiological. Several historical studies based on ion-selective microelectrode technique suggested the intracellular chloride concentration to be 10-20 mM (reviewed by Duan (2013)). However, the representativity of these experiments is limited, given the artificial experimental setting and the fact the cells were quiescent (e.g., markedly changing the role of I_(Ca)_Cl, reducing chloride influx to the cell when calcium transients are absent). The importance of experimental conditions on the variability of measured intracellular chloride concentration is also evident from the study in rabbit myocytes by Fong and Hinke (1981), where the measured chloride activity was 15.2–15.4 mM using HCO_3_–CO_2_ Ringer buffering, but 20 or 24.1 mM was observed when using HEPES-buffered Ringer, depending on the pH. A more recent study based on fluorescence dye measurements in murine myocytes at physiological temperature reported the intracellular chloride concentration to be ca. 29 ± 5 mM, with which our model is fully consistent. However, given the differences between human and murine physiology, the value is not conclusive for human. Therefore, more research is needed to clarify the physiological levels of intracellular chloride in human cardiomyocytes. Importantly, as shown in the next section, the simulations employing ToR-ORd-dynCl are in agreement with corresponding human experimental data, which is why we believe that the intracellular chloride concentration of 29.2 mM is not particularly problematic.

In summary, ToR-ORd-dynCl is capable of reaching a steady-state (using the term slightly loosely, in the sense of “without showing a visible drift”), whereas the original ToR-ORd shows a drift during long-term simulations as expected from the fixed intracellular chloride concentration (Hund et al., 2001). However, even for ToR-ORd, we note that simulations commonly employed in ventricular electrophysiology (of the order of seconds or minutes) correspond to the first few points in Figure 2), showing no substantial changes in key features or state variables. All diastolic ionic concentrations, resting membrane potential, action potential duration, and the state variable determining CaMKII activity changed by less than 1% during the first 1000 beats of the simulations (more than 15 minutes of simulated time).

In the rest of this document, the initial state for simulations of ToR-ORd-dynCl was set as the last recorded state after the 100,000 beats simulated here. I.e., all further results are in steady-state, not measuring the progress during the initial stabilisation. The initial state for simulations of the original ToR-ORd was kept as in the original study.

### 3.2 Performance of ToR-ORd-dynCl on single-cell calibration and validation criteria of ToR-ORd

To investigate the similarities and differences in the behaviour of ToR-ORd-dynCl and the original ToR-ORd, we re-ran the model evaluation pipeline described in the original article (Tomek et al., 2019) on the ToR-ORd-dynCl model. The pipeline runs the single-cell calibration and validation criteria summarised in Table 1 of the original study. We note that only two lines of code had to be changed in the script running the evaluation pipeline (which model function to use and which starting state to use) to have the evaluator run with the new model, demonstrating how easy to use the tool is, given the flexible code structure.

#### 3.2.1 Calibration criteria

Action potential and calcium transient are very similar between ToR-ORd-dynCl and the original ToR-ORd after 1000 beats of 1 Hz pacing, with less than 1 ms difference in APD90 and ca. 2 nM difference in calcium transient amplitude (Figure 3A,B). This means that the introduction of dynamic chloride concentrations does not markedly alter the behaviour of the model in the two most crucial features at the point when it was evaluated throughout the original article (after 1000 beats). The advantage of ToR-ORd-dynCl is that it maintains the action potential and calcium transient morphology even over extremely long simulations, as it is in the steady state.

**Figure 3:**
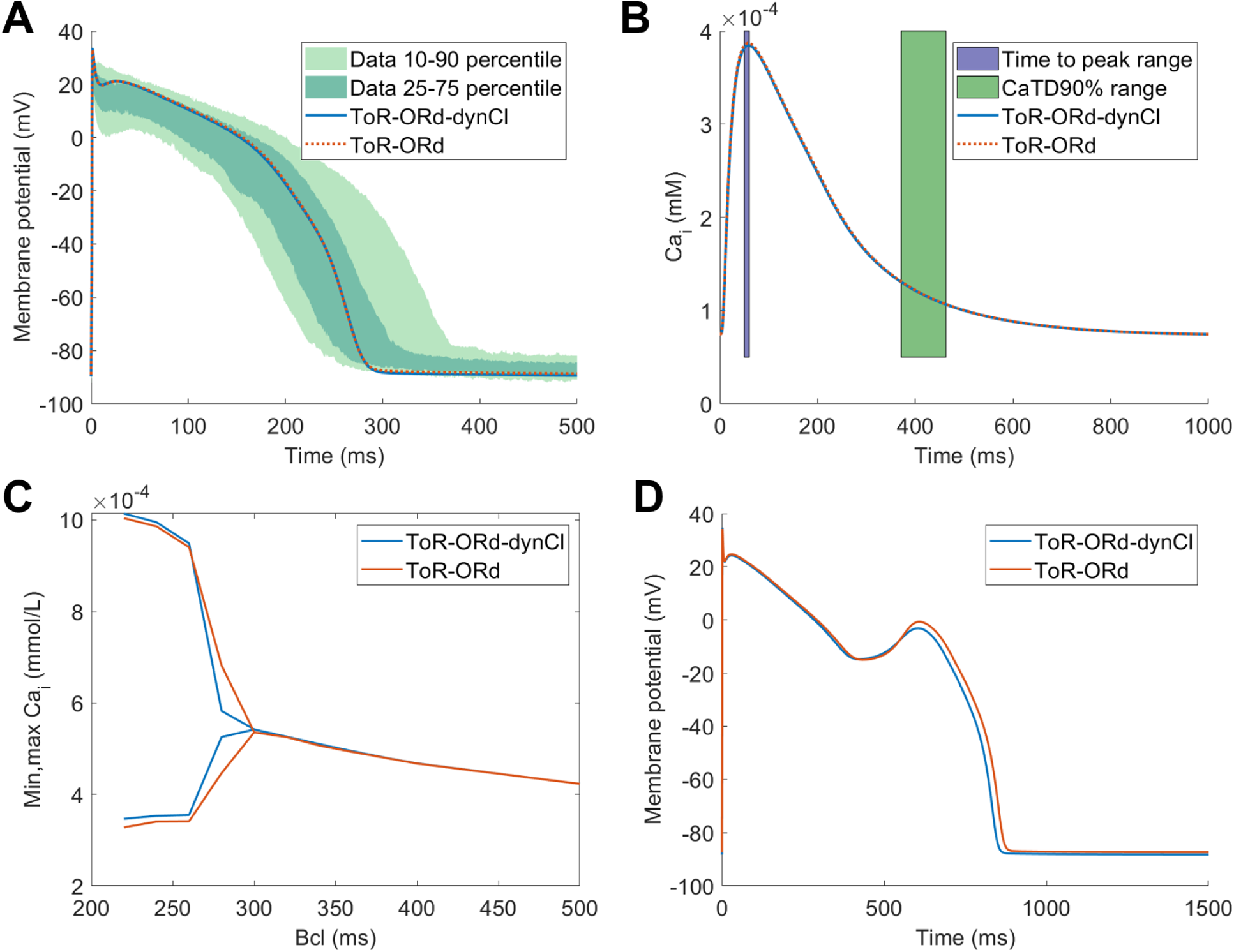
Calibration criteria and the comparison of ToR-ORd-dynCl with ToR-ORd. A) Action potential morphology, B) calcium transient, C) minimum and maximum calcium transient amplitude over a pair of consecutive 2 beats at a given basic cycle length; bifurcations indicate alternans, D) early afterdepolarisation formation at 0.25 Hz pacing with 85% block of I_Kr_ (with extracellular sodium of 137 mM and extracellular calcium of 2 mM, consistent with the reference experimental study Guo et al. (2011)). The dotted line style in A,B was used purely to allow plotting of near-identical curves and does not carry any additional meaning.

Alternans shows a similar pattern between the two models, being present at the same range of frequencies, with somewhat lower-amplitude alternans only at the basic cycle length of 280 ms in ToR-ORd-dynCl (Figure 3C). Both models manifest early afterdepolarisations of similar morphology (Figure 3D).

In addition, a sodium blockade (50% I_Na_+50% I_NaL_) reduced calcium transient amplitude by 7.14% in ToR-ORd-dynCl, which is similar to 6.19% in ToR-ORd. Negative inotropy of sodium blockade is well established (Bhattacharyya and Vassalle, 1982; Gottlieb et al., 1990; Legrand et al., 1983; Tucker et al., 1982), and it is important that ToR-ORd-dynCl maintains this property of ToR-ORd. Finally, a 50% I_K1_ blockade depolarises ToR-ORd-dynCl cell (from -89.75 mV to 89.03 mV), in line with ToR-ORd (from 89.06 mV to 88.44 mV) and with experimental literature (Dhamoon and Jalife, 2005).

#### 3.2.2 Validation criteria

ToR-ORd-dynCl gives similar results as ToR-ORd when simulating the effects of a range of channel blockers on action potential duration, and both models are well within the standard deviation from mean of the data (Figure 4A-D). Both models are within several milliseconds from each other when simulating APD accommodation to changes in pacing rate (Figure 4E). Ultimately, we simulated the S1-S2 restitution protocol in single cell, with both models giving near-identical results (Figure 4F).

**Figure 4:**
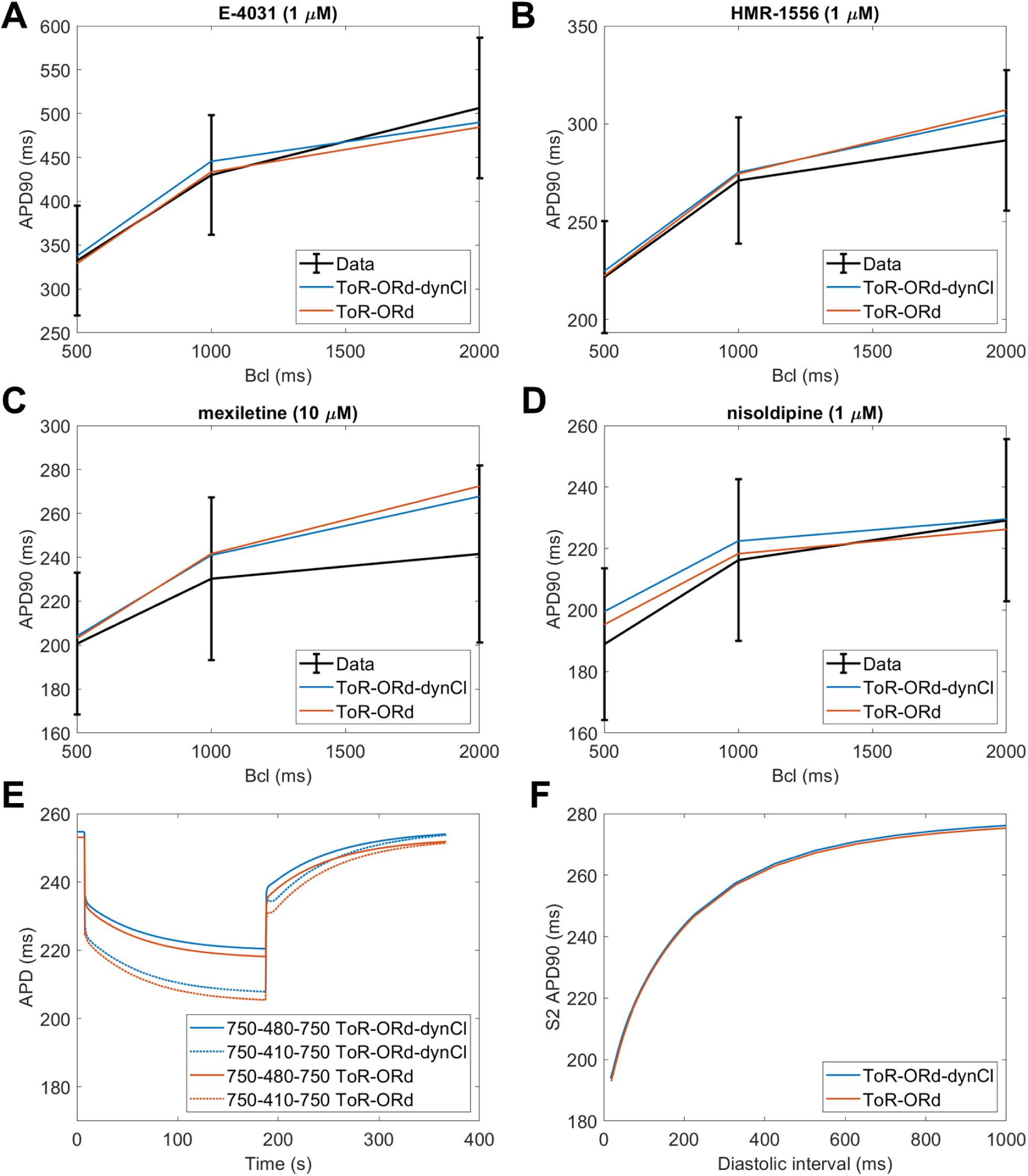
Validation criteria and the comparison of ToR-ORd-dynCl with ToR-ORd. A-D) The comparison of the two models and experimental data (based on O’Hara et al. (2011)) for four simulated compounds (see Tomek et al. (2019) for details): I_Kr_ blocker E-4031, I_Ks_ blocker HMR-1556, multichannel blocker mexiletine, and I_CaL_ blocker nisoldipine. E) APD accommodation for the two models. For both models, the baseline basic cycle length of 750 ms was changed to 410 or 480 ms before returning to 750 ms. F) Comparison of the S1-S2 curves in single cell (S1 = 1000 ms).

## 4 Conclusions

The ToR-ORd-dynCl behaves very similarly to the previously published ToR-ORd model in all observed features, gaining stability over time, while reaching a slightly higher chloride concentration and very slightly increased model complexity. Further experimental studies are required to determine the physiological intracellular chloride concentration in human cardiomyocytes.

## Acknowledgment

We thank Yann-Stanislas Barral for highlighting the issue of model stability and helpful discussion on this topic.

